# MIN1PIPE: A Miniscope 1-photon-based Calcium Imaging Signal Extraction Pipeline

**DOI:** 10.1101/311548

**Authors:** Jinghao Lu, Chunyuan Li, Jonnathan Singh-Alvarado, Zhe Charles Zhou, Flavio Fröhlich, Richard Mooney, Fan Wang

## Abstract

In vivo calcium imaging using 1-photon based miniscope and microendoscopic lens enables studies of neural activities in freely behaving animals. However, the high and fluctuating background, the inevitable movements and distortions of imaging field, and the extensive spatial overlaps of fluorescent signals emitted from imaged neurons inherent in this 1-photon imaging method present major challenges for extracting neuronal signals reliably and automatically from the raw imaging data. Here we develop a unifying algorithm called MINiscope 1-photon imaging PIPEline (MIN1PIPE) that contains several standalone modules and can handle a wide range of imaging conditions and qualities with minimal parameter tuning, and automatically and accurately isolate spatially localized neural signals. We quantitatively compare MIN1PIPE with other existing partial methods using both synthetic and real datasets obtained from different animal models, and show that MIN1PIPE has a superior performance both in terms of efficiency and precision in analyzing noisy miniscope calcium imaging data.

## INTRODUCTION

In vivo calcium imaging of activities from large populations of neurons in awake and behaving animals has become one of the staple technologies in neuroscience (Cai et al., 2016; Flusberg et al., 2008; Ghosh et al., 2011). Recent advances in single-photon based miniscope technology have further enabled imaging of neural ensemble activities in freely moving animals (Cai et al., 2016; Flusberg et al., 2008; Ghosh et al., 2011), thereby allowing circuits involved in a rich repertoire of animal behaviors to be examined. For example, this technology has been successfully used in probing dynamics of neural circuits involved in innate behaviors (Betley et al., 2015; Douglass et al., 2017; Jennings et al., 2015), decision making (Pinto and Dan, 2015; Poyraz et al., 2016), motor control (Klaus et al., 2017), learning and memory (Grewe et al., 2017; Kamigaki and Dan, 2017; Kitamura et al., 2017; Roberts et al., 2017; Roy et al., 2017; Xu et al., 2016), social memory (Okuyama et al., 2016), hippocampal place coding (Ziv et al., 2013), sleep (Cox et al., 2016; Weber and Dan, 2016), bird song (Markowitz et al., 2015), and pathological processes (Berdyyeva et al., 2016).

The increasing popularity of the miniscope calcium imaging technology demands the development of a fully automatic and robust signal processing method that can reliably extract neuronal signals from the noisy single photon calcium imaging data. Ideally, the processing method 1) should be able to handle a wide range of imaging conditions (*e.g*. high fluctuating background) and results with minimal parameter tuning, and 2) should have minimal assumptions about the quality of the data, such as free of movement or distortion, or sufficiently good signal-to-noise ratio (SNR). Existing imaging processing algorithms do not meet these two criteria.

For extraction of neuronal signals, many previous methods work well in situations with high SNR and stable field of views, therefore they are best suited for processing two-photon imaging data. These algorithms include linear unsupervised basis learning methods (Mukamel et al., 2009; Reidl et al., 2007; Pachitariu et al., 2013), nonlinear unsupervised methods (Maruyama et al., 2014; Pnevmatikakis et al., 2016), and the supervised learning method (Apthorpe et al., 2016). The PCA/ICA (principal component analysis followed by independent component analysis) method (Mukamel et al., 2009) was the first attempt to automatize the signal extraction process from miniscope imaging data through manual annotations of ROIs, but this method has difficulties in delineating localized ROIs and in separating overlapping ROIs. Since single-photon based imaging collect lights from a large depth of field, overlapping neurons in different depth is not uncommon. Another method called CNMF (constraint matrix factorization framework) (Pnevmatikakis et al., 2016), which combines nonlinearity in matrix factorization with simultaneous deconvolving spike trains from calcium dynamics, returns more spatially localized maps of ROIs compared to other methods and has better performance in identifying overlapping neurons. While methods like CNMF achieve plausible results in processing two-photon imaging data, they are ill-suited for processing the single-photon based miniscope imaging because: 1) the data from miniscope imaging are dominated by noisy, uneven and fluctuating background, 2) such methods depend on sophisticated parameter tuning, especially requiring setting parameters that are unknown *a priori* in practice such as “number of neurons”. Thus, a new method that can effectively remove background yet preserve real neural signals is highly desired.

Moreover, both PCA/ICA and CNMF rely on the stable imaging field and will fail if the imaging field contains movement, but movements (including distortions) during imaging, are often inevitable. Therefore, correcting movements and distortions is another major problem needs solving before neural activity signals can be reliably extracted. Many methods were developed independently in attempt to solve this problem. For example, several approaches register frames through template matching based on the assumption that the major form of movements is translational displacement (Dubbs et al., 2016; Thévenaz et al., 1998). To eliminate such assumptions, methods with block-based displacement field estimation were developed, with image feature matching algorithms extended from Lucas-Kanade tracker (Greenberg and Kerr, 2009; Lucas and Kanade, 1981) or Hidden Markov Models (Dombeck et al., 2007; Kaifosh et al., 2013). Some of these movement correction methods have been included as a module in a larger toolbox, using such frame-wise rigid registration approaches (Kaifosh et al., 2014; Pachitariu et al., 2017). When applied to handle nonrigid movement registrations, existing methods make specific assumptions about both the form and magnitude of the potential movements that result in suboptimal performance when large deformation occurs during imaging. Furthermore, errors in the movement correction can easily propagate since these methods all register frames based on a single reference frame. Considering that extracting neural activity signals is highly dependent on removing movement artifacts, an accurate and robust movement correction module is imperative. A simple unimodal algorithm for either translation/rigid or nonrigid registration is insufficient for this purpose.

Here we develop the MINiscope 1-photon imaging signal extraction PIPEline (MIN1PIPE) that takes the very raw calcium videos as inputs, and automatically removes background while preserving signals, corrects movements with no assumptions of the types of movements, and delivers separated neuronal ROIs as well as deconvolved calcium traces as outputs. The MIN1PIPE contains a neural enhancing module that minimizes the influence of background unevenness and fluctuations, a hierarchical movement correction module that can handle all kinds of deformation with minimal error propagation, and a seeds-cleansed neural signal extraction module that identifies the set of real ROIs and their corresponding calcium traces without setting unknown parameters *a priori* (Fig. 1). Though our MIN1PIPE is primarily developed for single-photon based miniscope imaging, individual modules can also be independently combined with other processing algorithms to improve performance in analyzing two-photon imaging.

**Fig. 1.**
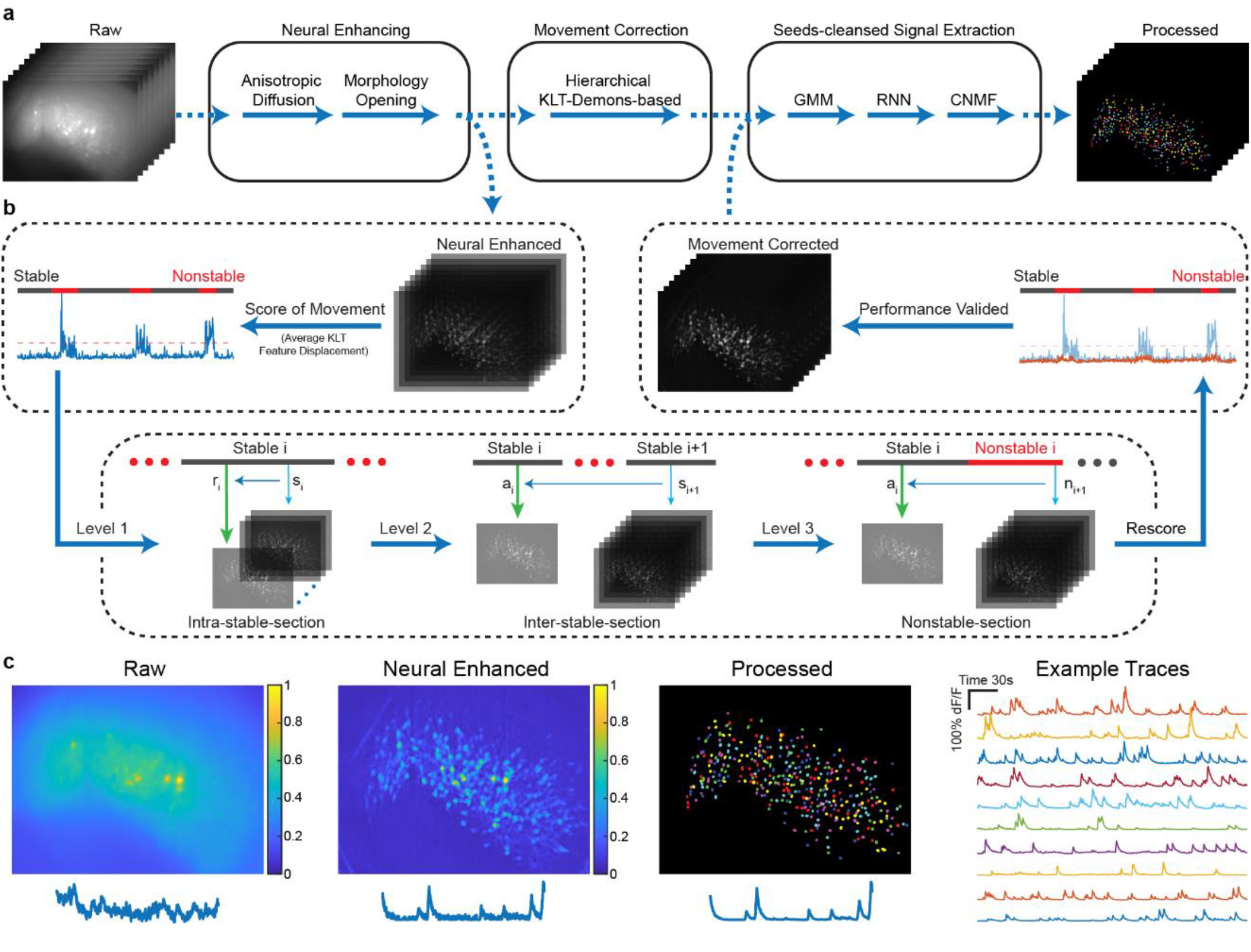
The general pipeline and demonstrations of MIN1PIPE. **a.** The overall structure of MIN1PIPE. MIN1PIPE takes the very raw miniscope imaging data freshly collected from the imaging system as inputs, and returns fully processed ROI components with spatial footprints and temporal calcium traces as outputs. The data are processed in series by neural enhancing, hierarchical movement correction, and seeds-cleansed signal extraction modules. Each module is composed of specific brick functions. **b.** A zoom-in of the hierarchical movement correction module. The module is KLT-Log Demons based. It first scores all the neural enhanced frames and divides them into stable and nonstable sections, then register frames at three different levels. The movement corrected frames are then fed into the seeds-cleansed neural signal extraction module. **c.** The max projecting demonstrations of MIN1PIPE. The raw imaging video contains large, dominant background fluctuation, the neural enhanced video contains mainly neural signals, and the fully processed video contains only extracted and denoised neural signals as independent ROI components. Example traces are randomly selected from the processed video. See also Fig. S1-S4.

## RESULTS

### Core modules in MINIPIPE

#### Neural Enhancing Module

The core of MIN1PIPE relies on turning the imaging data into a stack of background free, baseline corrected frames as the first step. Due to the complex spatiotemporal properties of background dynamics, our method applies a frame-wise background estimator that is adaptive to the local properties. A natural idea can be translated from mathematical morphology and computer vision, that the foreground neural signals can be approximated by subtracting the estimated background in a denoised image. Therefore, we first remove the grainy noise inherent to the single-photon system (see example of such grainy noise in Supplementary Fig. S2) while preserving the boundary between foreground and background, and this is achieved by applying an *anisotropic diffusion* denoising operation (Perona and Malik, 1990) on the raw imaging frames. Next, we use a simple straightforward *morphological opening* operation (Serra and Vincent, 1992) as the background estimator, with the size of the structure element similar to that of the neurons in the imaging field. The opening operation removes structures smaller than the desired structure element. Subsequently, the foreground that contains all the neuronal signals with the minimal noise is computed as the difference between the denoised raw and the morphological opened frames (Fig. 1a and Supplementary Fig. S1-S2).

#### Movement Correction Module

After the neural signals are enhanced, we next correct for movements in the imaging videos. The problem of movement correction can essentially be broken down to image stack registration. However, without setting specific constraints on the form or magnitude of the movements, even the most efficient registration algorithms require a running time on the order of seconds to minutes per frame (Vercauteren et al., 2009). Considering that the general imaging datasets contain tens of thousands of frames, the time required for applying these sophisticated image registration methods to every frame is inconceivable. Here we develop a hierarchical video registration framework for the correction of all types of movement, without sacrificing the precision or the speed of corrections. Our framework first decomposes the imaging video into two sections: the *stable sections* whose movements can be approximated by small translational displacement, and the *non-stable sections* that contain large general deformation. This step uses the KLT tracker that estimates the displacement of potential corner-like features between two neighboring frames (Shi and Tomasi, 1994). Next, we employ three levels of different strategies to align images. At the first level, we correct the small translational displacement within each stable section using the fast Lucas-Kanade tracker, which can be performed efficiently in parallel on multiple sections (Lucas et al., 1981). We then incorporate a diffeomorphic Log-Demons image registration method which can handle large deformations while preserving the local geometrical properties (Vercauteren et al., 2009). At the second level, we align all the stable sections. The overall information of each section is extracted to form a sectional image. The current sectional image is then aligned to a reference sectional image, which is generated as a linear summation of all previously registered sectional images that is the closest to the current image. The summation weights are determined by least square regression between the current and previous sectional images. The estimated displacement field is then applied to each frame within the current section. This will be iterated until all stable sections are aligned. At the third level, we use a similar process to register the individual frames within each non-stable section, which can be parallelized to boost performance efficiency (Fig. 1b). This hierarchical approach significantly reduces the total registration time due to the balanced assignment of different methods. Importantly, the common registration error does not propagate with this approach.

#### Seeds-Cleansed Neural Signal Extraction Module

Once movements are corrected and images are aligned, the main task turns to the neural signal extraction. MIN1PIPE extracts neural signals automatically in two main steps, 1) the seeds cleansing step to reliably detect the set of real ROIs, and 2) an altered spatiotemporal CNMF to separate ROIs and corresponding calcium traces (Pnevmatikakis et al., 2016). Previous methods contain either no explicit seeds initialization step or only a coarse initialization that compromises between precision and recall. In contrast, our seeds cleansing step forces the algorithm to find the set of real ROIs. This is achieved by first generating an over-complete set of seeds containing all potential centers of real ROIs at the cost of including false positives. This over-complete set is then coarsely cleansed by applying a two-component Gaussian Mixture Model (GMM) on the peak-valley difference of corresponding traces of the seeds, where the traces of real neurons usually have larger fluctuations compared to the nonneuron false positive seeds. The GMM removes most background false positives without losing real neurons (Supplementary Fig. S3). To further remove the remaining false positives, such as the ones with abnormal background fluctuations or hemodynamics, we have trained an Recurrent Neural Networks (RNNs) with LSTM module offline as the classifier for calcium spikes (Supplementary Fig. S4) (LeCun et al., 2015; Hochreiter and Schmidhuber, 1997). Those seeds whose traces contain RNN-identified calcium spikes, regardless of their temporal locations, are classified as true positives whereas the rest are deemed as false positives. After such cleansing processes, there is still a low possibility of identifying multiple seeds within a single ROI. Therefore, we merge potentially redundant seeds by computing the temporal similarity of seeds within their neighborhoods, and preserving the ones with maximum intensity. With the cleansed set of seeds as the initial position of ROIs, we next perform the iterative spatial and temporal optimizations, as proposed in CNMF, to update the spatial footprints of individual ROIs, and the temporal traces with deconvolved spike trains. Notably, unlike previous CNMF, where the spatial footprints are sequentially updated and subtracted from the preceding residuals, we extract spatial footprints from the original data that does not depend on preceding iterations. Therefore, the information loss/duplication is reduced and the optimization procedures can be parallelized in our method.

We show example results obtained using the MIN1PIPE methodology including a raw frame, the fully processed ROIs and the example traces from ROIs (Fig. 1c).

### Quantitative validation of the MIN1PIPE performance

The neural signal extraction and movement correction are two independent problems that can be tested separately. For the signal extraction, we test the performance on synthetic datasets with various signal levels, while for the movement correction, we can directly test it on real data. To measure the performance of the signal extraction, we use a scoring metric that evaluates the spatial and temporal similarity between the ground truth and the identified ROIs (see Online Methods), and calculate true positive, false positive and false negative. To measure the performance of the movement correction, we use a metric based on the average displacement of feature points between neighboring frames.

We synthesized 16 imaging videos with signal levels (SL, defined as the ratio of the amplitude between the signals and the background) ranging from 0.05 to 0.8. Each video contains 3000 frames with 100 neurons of various shapes and calcium dynamics, and background fluctuations extracted from real datasets (details in Supplementary Notes S1-S2). The properties of synthetic videos resemble those of real data, whose SL falls between the range of 0.2 and 0.8. The condition at 0.05 SL is an extreme (Supplementary Fig. S5 and Video S1-S3). We compare MIN1PIPE with PCA/ICA and CNMF. We used commercially available *Mosaic* software (Inscopix Inc.) which implements PCA/ICA method (Mukamel et al., 2009), and processed the data following the standard workflow in the software manual. In particular, we chose the number of principal components (PC) and independent components (IC) based on the suggested rate (e.g. 20% more ICs and 50% more PCs than the estimated number of ROIs). For CNMF, we used the default initialization strategy in (Pnevmatikakis et al., 2016).

The results of ROI detection (Fig. 2a) and the calcium traces from one example ROI obtained by different methods (Fig. 2b) at 0.2 and 0.8 SL are compared. The contours of the identified ROIs using different methods are drawn and superimposed onto the max projection of the ground truth. For both SLs, MIN1PIPE can identify the nearly complete set of ROIs (93% and 100% for 0.2 and 0.8 SL respectively) with minimal false positives (1% and 0% respectively), whereas the other two methods detect partial subsets of ROIs (PCA/ICA: 0% at 0.2 SL and ~65% true positives at 0.8 SL; CNMF: ~32% at 0.2 SL and ~95% true positives at 0.8 SL). In addition, the extracted spatial footprints are less realistic with the PCA/ICA or previous CNMF methods. The example calcium traces indicate that MIN1PIPE has a near-optimal performance in extracting individual ROIs even at low SLs when compared to the ground truth traces (Fig. 2b). In contrast, PCA/ICA completely fails to identify ROIs when SL is low, while CNMF fails to separate neurons from overlapping ROIs. Fig. 2c summarizes the performance accuracy of these three methods at all SLs. The plots of the true positive, false positive and false negative indicate that MIN1PIPE has a significantly superior performance in all conditions, and outperforms the other two methods in the extreme conditions. Fig. 2d summarizes the spatiotemporal similarities between the extracted results and the ground truth. The clusters of point clouds confirm again that MIN1PIPE outperforms the other two methods in best resembling both the spatial and temporal properties of the ground truth signals. The results for all signal levels are shown in Supplementary Fig. S6.

**Fig. 2.**
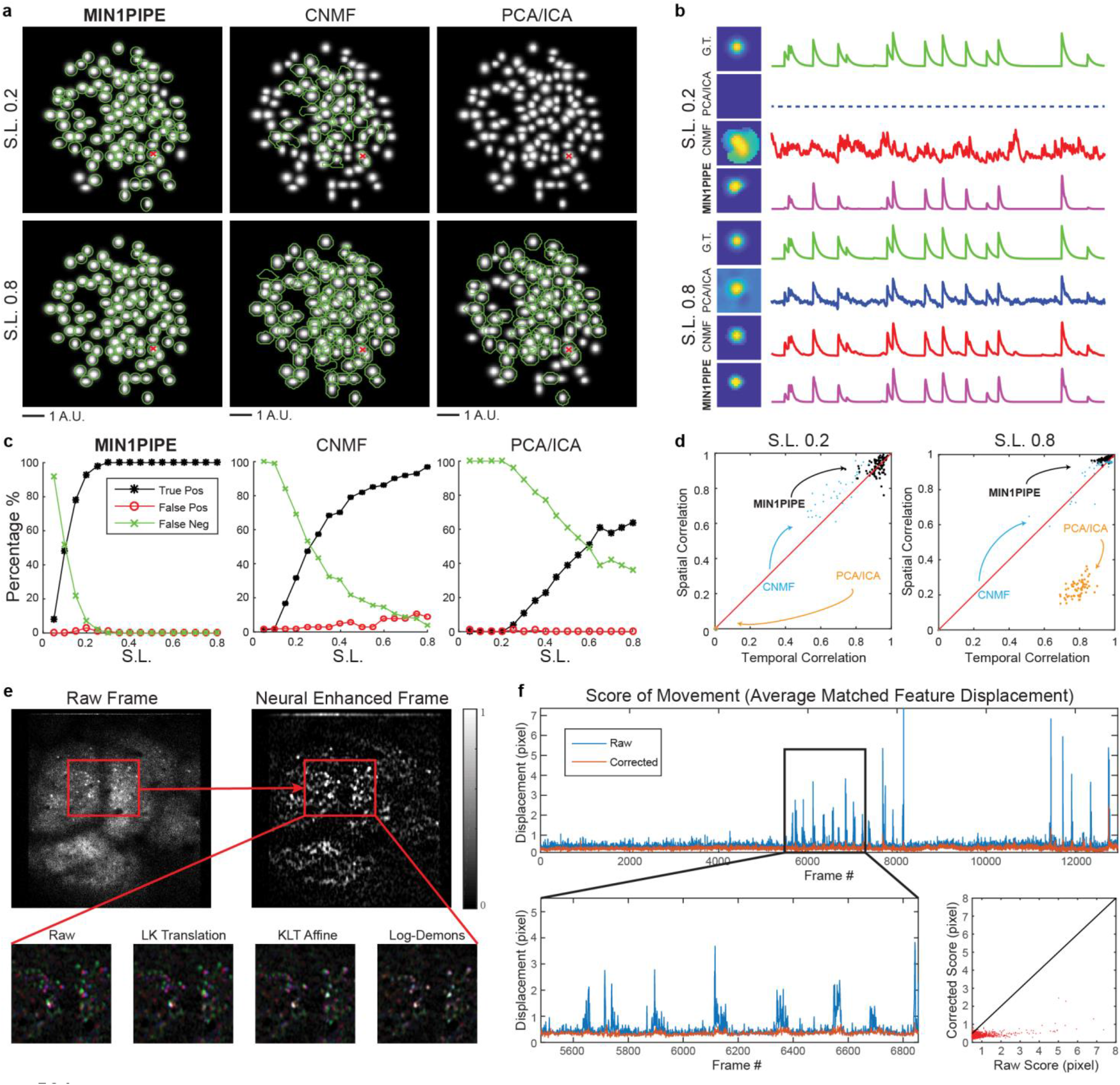
Quantification of MIN1PIPE. **a-d.** The quantitative performance comparison between MIN1PIPE and the other methods on simulated datasets. **a.** The identified ROI contours of the three methods at two representative signal levels. S.L.: signal level (the ratio of the amplitude between the signals and the background). A.U.: arbitrary unit. **b.** The trace of an example ROI component identified by the three methods and the ground truth. G.T.: ground truth. **c.** The identification precision and accuracy of the three methods. **d.** The spatiotemporal identification accuracy with the three methods at two representative signal levels. The similarity of the spatial footprints and temporal traces between the results of the three methods and the ground truth of each ROI is plotted as a dot in the figure. These quantifications confirm that MIN1PIPE not only outperforms the other two methods in all aspects, but also verify that MIN1PIPE can provide satisfactory neural signal extraction results of miniscope imaging data. **e-f.** The quantification of the movement correction module. **e.** The demonstration of various image registration methods in a consecutive three frames from two-photon imaging data. The deformation registration algorithm we use in the module (Log-Demons) successfully corrects all the nonrigid deformations within the frames that cannot be corrected by the methods that assume translation or rigid transformation. The black and white color indicate overlapping of the same structure between the frames, whereas other colors indicate nonaligned structures. **f.** The score of movement before and after the correction. In general, the hierarchical correction steps register large deformations while preserves the stable frames. See also Fig. S5-S9.

To validate the performance of the movement correction in MIN1PIPE, we applied the module on video sections with large deformation movements of the imaging field. Specifically, we chose a video obtained through two-photon imaging of the ferret’s posterior parietal cortex as an example due to its particularly frequent and large deformations (Supplementary Video S4 as a demo section of the full video). Fig. 2e shows an example of 3 consecutive frames (with each frame pseudo-colored as pink, green or blue) with large nonlinear deformations between frames superimposed together. The two rigid-transform-based methods (LK and KLT Affine) by themselves fail to fully remove the large deformations, whereas MIN1PIPE (combining the Log-Demons transformation) succeeds in correcting these movements. We further quantified the extent of correction by calculating the average displacement of feature points between two neighboring frames before and after the movement correction (Fig. 2f). Before correction, the score of average displacement shows frequent burst of large deformation periods, whereas after correction, the score of displacement shows a nearly flat line. Notably, the frames with large deformations are all well aligned (Fig. 2f lower panels; Supplementary Video S4).

### Application of MINIPIPE on real miniscope imaging datasets

We next compare the performance of MIN1PIPE with PCA/ICA and CNMF methods on real datasets. We first applied the three methods to the miniscope calcium imaging data obtained using prism probe from layer 2 and 3 of the barrel cortex in freely moving mice. GCaMP6f was expressed in layer 2/3 neurons using AAV, and signals were imaged continuously over 5min when the mouse freely explored its environment. Following the general pipeline of MIN1PIPE (Online Methods), our method removed the strong uneven background structure, and automatically identified 210 putative ROI components without the need for additional manual selection (Fig. 3a, Supplementary Video S5). In comparison, PCA/ICA identified 79 ROIs whereas CNMF identified 71 ROIs. Note that with the same computer configuration and dataset (~28 GB), the CNMF ran into memory issues and could not process the full-scale video, thus we cropped a center patch as the input video to the CNMF (Fig 3b-d middle panels). The contours of the ROIs identified by the different methods are drawn and superimposed onto the max projection of the neural enhanced data (Fig. 3b). This reveals that MIN1PIPE can identify potentially all ROIs, whereas PCA/ICA and CNMF miss a significant subset of ROIs. Meanwhile, both the PCA/ICA and CNMF have some problems in separating overlapping ROIs and/or estimating the correct shape of the ROIs, as revealed by the max projections of the extracted signals (Fig. 3c). The projection of MIN1PIPE extracted signals closely resembles those of the neural enhanced data (Fig. 3c, top). In comparison, the projection obtained using PCA/ICA has low SNR with high background signals (Fig. 3c, bottom), whereas the projection derived from CNMF shows unrealistic ROI shapes larger than the true shape of neurons, indicating that the CNMF is sensitive to the contamination of the background dynamics (Fig. 3c, middle). To further check the shape of individual ROIs, we choose to visualize one example ROI embedded in the full imaging field using different methods (Fig. 3d). MIN1PIPE delineates a well localized ROI footprint, whereas CNMF returns a less localized ROI difficult to relate to the underlying neuron. PCA/ICA, on the other hand, fails to provide a localized footprint as remnants of other ROI components can also be seen. To examine the temporal traces extracted by the different methods, we selected 10 ROI components that were identified by all three methods within the cropped image field and plotted their corresponding calcium traces (Fig 3e, individual panels marked with 1-10). Again, the spatial footprints of the 10 ROIs obtained through CNMF and PCA/ICA are less localized as described above. In terms of calcium dynamics, both MIN1PIPE and CNMF give denoised traces, whereas the traces obtained using PCA/ICA are noisy and show unrealistic negative fluctuations. Furthermore, the traces extracted using CNMF include false positive calcium events that likely reflect background noises (arrows in Fig. 3e). Supplementary Video S6 shows the comparison of raw and processed results using three different methods of the entire imaging video.

**Fig. 3.**
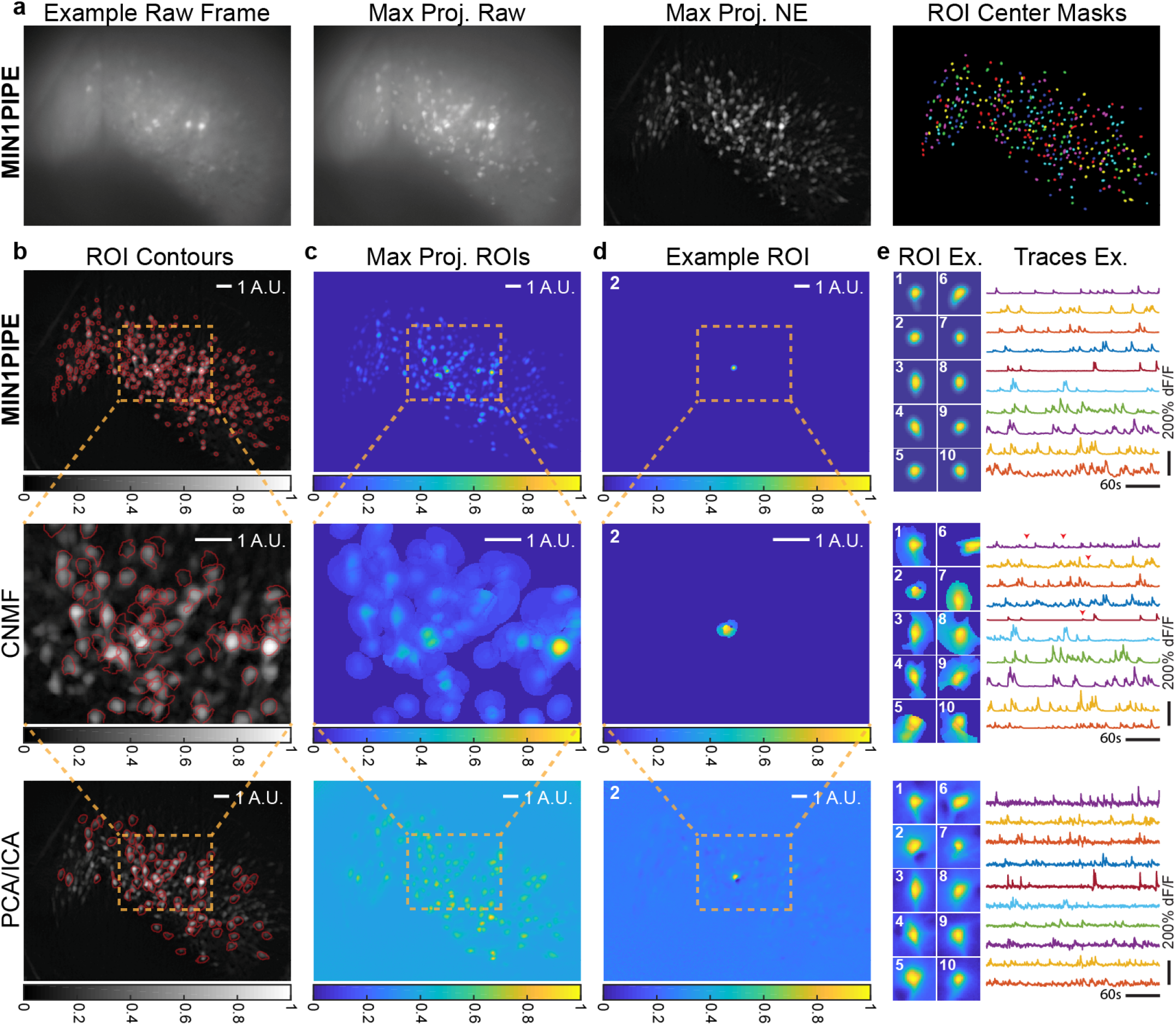
Comparing different methods using miniscope imaging data from the mouse barrel cortex. **a.** Demo of MIN1PIPE in processing imaging data from the barrel cortex. We show an example raw frame, the max projection of raw video, the max projection of the neural enhanced video, and the colored ROI masks. **b.** The identified ROI contours superimposed on the max projection of neural enhanced video. MIN1PIPE returns a set of ROIs with well-shaped, localized spatial footprints. CNMF and PCA/ICA identify only a small subset of true positives (71 and 79 respectively), and either the footprints are not localized, or the run into memory issues. The projection map is shown in grayscale colormap. **c.** The max projections of all identified ROI footprints. This further demonstrates the general properties of the extracted neural components. **d.** The spatial footprint of an example ROI. MIN1PIPE extracted the most localized component that is close to the real shape shown in the neural enhanced projection, whereas CNMF detected the noise-sensitive result and PCA/ICA detected the unrealistic component as a mix of several other components. **e.** The temporal traces of ten examples randomly selected from the ROIs that are detected by all three methods. A similar conclusion can be drawn in spatial footprints, whereas MIN1PIPE shows the best performance in temporal trace extraction (some potential false positive events indicated by red arrows).

To further illustrate the general applicability of MIN1PIPE in processing miniscope imaging data obtained over different brain areas and/or different animal models, we applied it to process calcium imaging results from Area X in zebra finch and compared the results with those obtained using the other two methods. Briefly, MIN1PIPE detected 55 ROI components, whereas CNMF detected 15 and PCA/ICA detected 35 ROIs after manual selection (Fig. 4a-b, Supplementary Video S7). Again, CNMF could only process a cropped portion of the imaging video due to the computer memory issues. All of the ROIs detected by CNMF and PCA/ICA are included in the set extracted by MIN1PIPE. In addition to false negatives, CNMF also identifies a cluster of false positive ROIs that are apparent upon visual inspections (Fig. 4b-c, e). The PCA/ICA again gives rise to nonlocalized ROI footprints that contain other potential components (Fig. 4d). It was known that Area X neurons show strong song selective activities when the bird is singing (Goldberg and Fee, 2010; Kojima and Doupe, 2007; Woolley et al., 2014; Yazaki-Sugiyama and Mooney, 2004), which can be used as partial ground truth to validate MIN1PIPE. We plot the calcium traces of the ROIs identified by MIN1PIPE, and the majority of the ROIs contain calcium events that are roughly phase-locked to the singing onsets (Fig. 4f). Notably, a subset of the neurons shows precise singing-related activities with minimal events unrelated to singing (Fig. 4g upper panel). Furthermore, we sorted neurons according to their timing of peak calcium activities during each song production event (Fig. 4g lower panel), and this analysis also reveals a subset of Area X neurons whose activation patterns are closely related to the onset of the singing, consistent with previous findings.

**Fig. 4.**
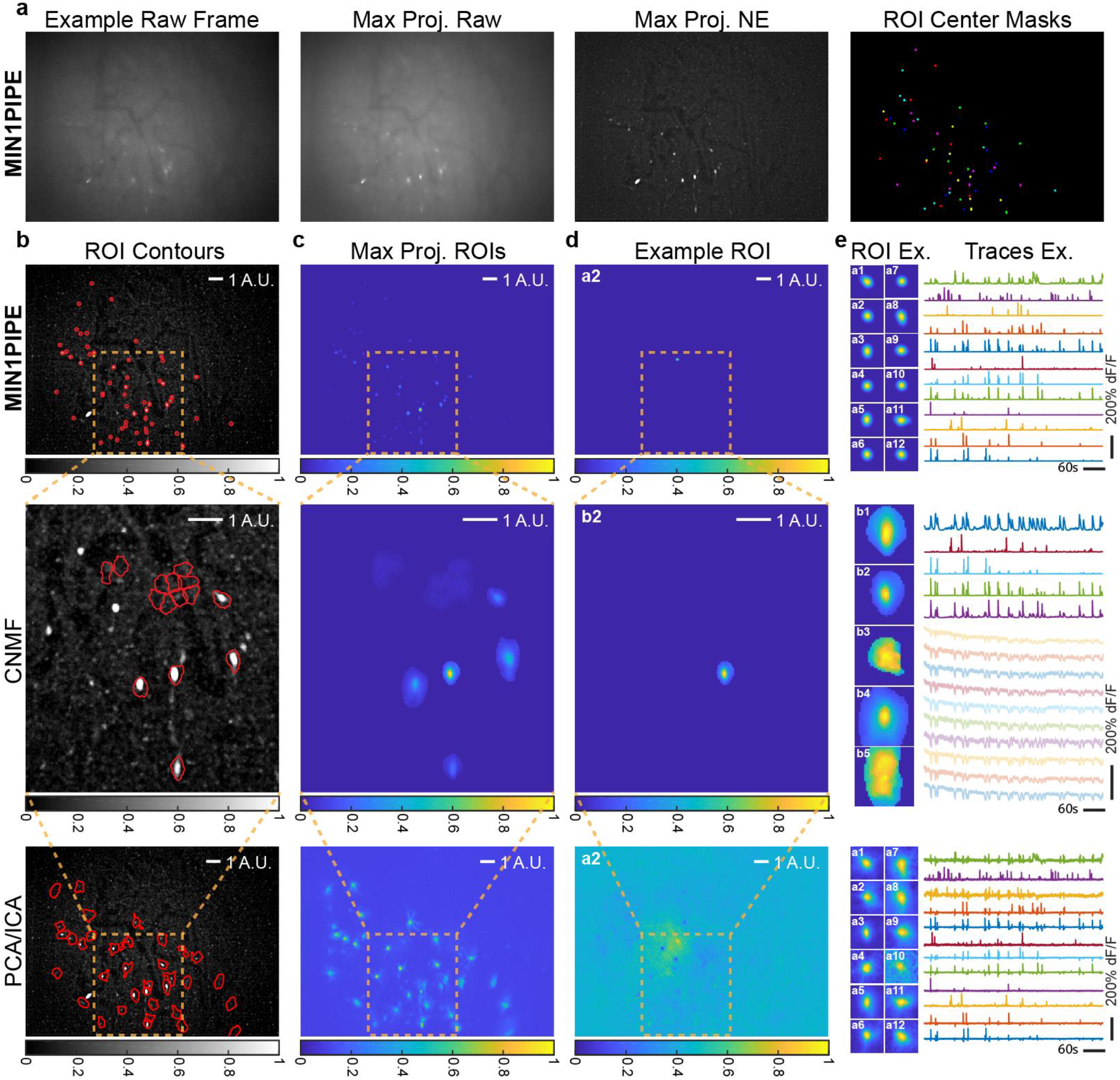
Comparing different methods using miniscope imaging data from the Area X in zebra finch. **a.** Demo of MIN1PIPE in processing data from Area X in the zebra finch brain. We show an example of the raw frame, the max projection of raw video, the max projection of the neural enhanced video, and the colored ROI masks. **b.** The identified ROI contours superimposed on the max projection of neural enhanced video. MIN1PIPE returns a set of ROIs with well-shaped, localized spatial footprints. CNMF and PCA/ICA identify only a small subset of true positives (15 and 35 respectively), and either the footprints are not localized, or the processes run into memory issues. The projection map is shown in grayscale colormap. **c.** The max projections of all the identified ROI footprints. **d.** The spatial footprint of an example ROI. MIN1PIPE extracted the most localized component that is close to the real shape shown in the neural enhanced projection, whereas PCA/ICA extracted the unrealistic component as a mix of several other components. CNMF on the other hand, did not detect this ROI. **e.** The temporal traces of twelve examples selected from the ROIs detected by both MIN1PIPE and PCA/ICA. A similar conclusion can be drawn in spatial footprints, whereas MIN1PIPE shows the best performance of temporal trace extraction. Faded traces in CNMF indicate the traces of the false positives.

We also did studies (Supplementary Fig. S7-S8) showing the effectiveness of the two modules (neural enhancing and seeds-cleansed signal extraction) when combined separately with previous methods to improve the performance. Finally, we compare our method with the CNMF-E method (Zhou et al., 2016) (Supplementary Fig. S9).

## DISCUSSION

The key advances that set MIN1PIPE apart from the previous imaging processing methods are the following. First, MIN1PIPE solves the full range of problems for signal extraction in single-photon miniscope imaging with one pipeline. Specifically, we have developed innovative and robust modules to solve different problems including: the neural enhancing module that uses a novel algorithm to remove the ultra-high and fluctuating background characteristic of single photon imaging, the novel hierarchical movement correction module that is capable of efficiently registering any types of deformations of the imaging field, and the seeds-cleansed neural signal extraction module that utilizes GMM and pretrained RNN to enable automatic identification and extraction of ROIs and calcium traces. Second, MIN1PIPE eliminates the need for heuristically setting many parameters that are not only unknown *a priori*, but also influence the performance of the downstream processing steps, such as setting the number of neurons, a central parameter required by all previous neural signal extraction methods. The importance of this should not be neglected, because pre-setting unknown parameters can become problematic in practice. For example, overestimating the number of neurons may result in false positives in identified ROIs before the calcium trace extraction step, and also result in the unnecessary consumption of computing time. These false positives can only be removed with laborious manual selection without a robust seeds-cleansing step. On the other hand, underestimating the number of neurons will likely lead to false negatives that can never be identified by the downstream steps. Therefore, tweaking this parameter is inevitable in practice using previous methods. While we do not claim that MIN1PIPE completely eliminates the need for manually pruning identified ROIs, our method does only involve minimal manual interference. Third, MIN1PIPE contains a minimal set of parameters that are easy to interpret and error-tolerant (Online Methods). These include a set of simplified and fixed parameters applicable to various conditions of the popular miniscope platforms (e.g. Inscopix nVista, UCLA open-source miniscope) in our brick algorithm. In addition, all modules use definitive criteria independent of various imaging datasets, which ensures the robustness that the previous combination of setting the number of neurons and the serial initialization procedure could not provide.

In summary, MIN1PIPE provides a generally applicable high-performance toolbox with the modular framework to handle and process miniscope imaging data. Interesting future works may integrate more advanced methods to further improve the precision, such as stricter choices of the kernel of anisotropic diffusion (Chen et al., 2011; Tsiotsios and Petrou, 2013), considering spectral invariants to handle very large deformations during non-rigid registration (Lombaert et al., 2014), and using more biologically valid calcium dynamics deconvolution methods (Speiser et al., 2017). Additionally, a more robust RNN classifier for seeds cleansing can be trained with more available datasets.

**Fig. 5.**
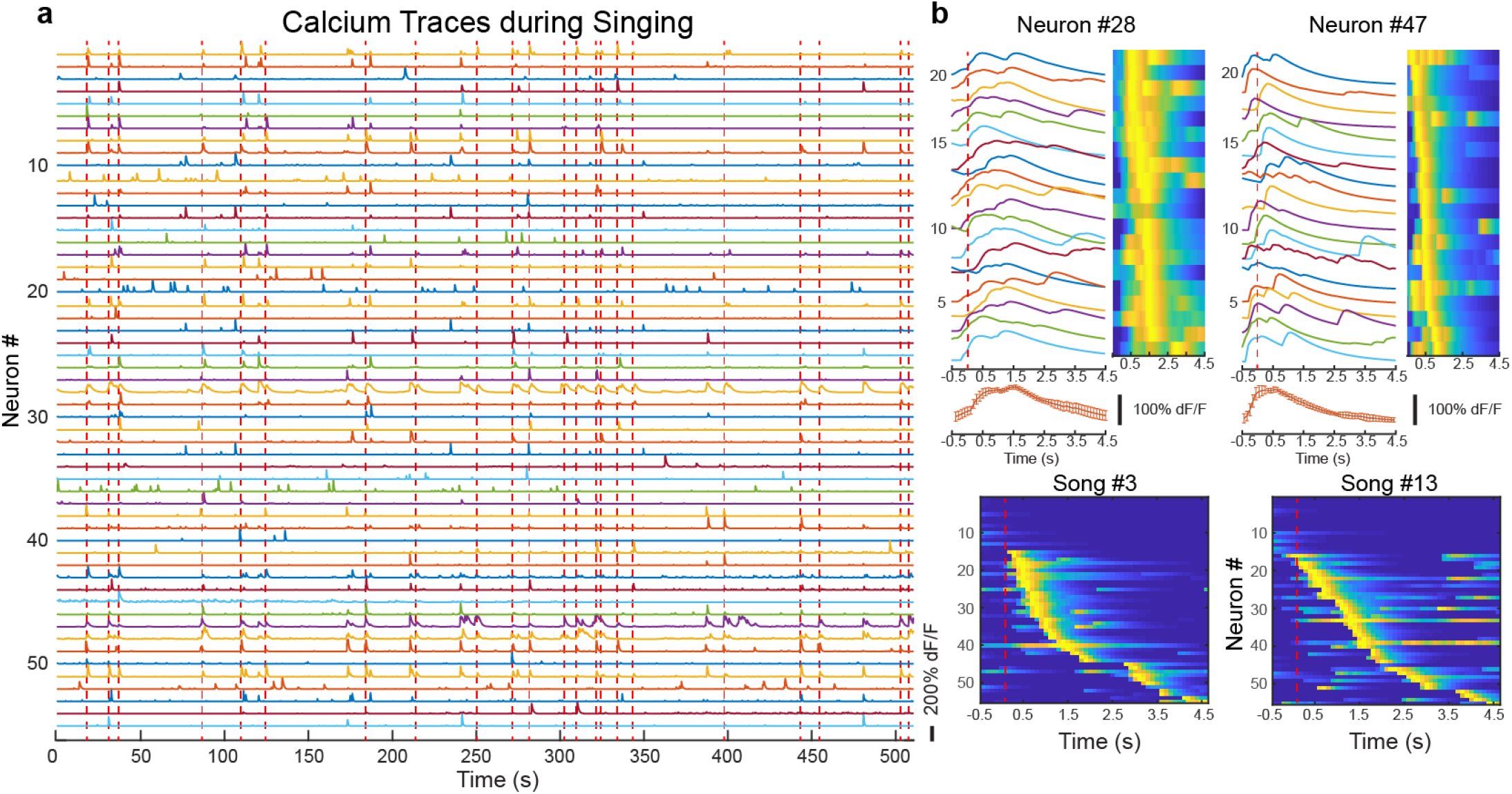
Analysis of neural correlation in Area X of zebra finch to singing behavior. **a.** The traces of all automatically detected ROIs with MIN1PIPE superimposed with complete singing onsets in red dashed lines. Note that incomplete song events are not labeled and taken into analysis. **b.** Upper panel: two example neurons with precise song selectivity. The traces represent the neuronal activity of the first 20 complete song singing events with a window of 0.5 second before and 4.5 seconds after the song onset. The heatmap is another way of visualizing the analysis. The error bar graphs at the bottom show the trial average traces of the two neurons. Lower panel: population activity pattern of two example song singing events. The neurons are sorted to the latency of the peak. Red dashed lines indicate the onset of songs.

## ACKNOWLEDGEMENT

We thank the suggestions and comments from members of the Wang Lab and Mooney Lab at Duke University. The study is supported by NIH grants DP1-MH103908, R21NS101441, NS077986 to F.W.

## AUTHOR CONTRIBUTIONS

J.L. and F.W. conceived and designed the project; J.L. designed and implemented all modules of MIN1PIPE; C.L. designed the RNN classifier; J.L. generated the synthetic data; J.L., F.W., J.S.-A., R.M., Z.Z. and F.F. designed and/or conducted in vivo imaging experiments; J.L. and C.L. analyzed the data; J.L., C.L., F.W. wrote the manuscript.

## ONLINE METHODS

### NEURAL ENHANCING

The neural enhancing module contains two steps of framewise operations. We denote 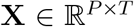 (*P* is number of pixels per frame, and *T* is number of frames) as the raw video after converting each two dimensional frame into a vector, and 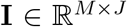 (*M* and *J* are the height and width of a frame respectively) as a frame before vectorization.

#### Denoising

To first remove the spatial noise embedded in each frame introduced by the photoelectric process of the sensor, we apply denoising operation on each frame **I** using anisotropic diffusion [36]. For a given diffusion time *τ*, the evolution follows the equation 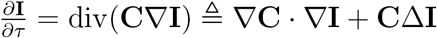 where **C** is the diffusive coefficient matrix depending on pixels and *τ*. By choosing concrete form of **C** and *τ*, we can control the smoothing level along or perpendicular to the boundaries between neurons and the background. Due to the simple structures in the cleaned imaging field, we choose the classical Perona-Malik filter 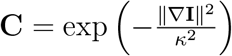, where *κ* controls the threshold of high-contrast. The diffusivity is selected to preferentially smooth high-contrast regions, and *κ* and *τ* are chosen to allow high tolerance. We use *κ* = 0.5 and *τ* = 0.5 with a step size *δt* = 0.05 as the default parameters in anisotropic diffusion. The output image **I^s^** contains reduced spatial noise while preserves the boundary information of ROIs.

#### Background Removal

Based on the observation that neuronal ROIs are small in size compared to background structures, the gray-scale morphological opening operator with a binary structuring element matrix **Φ** can well estimate the background B from the frame I adaptively. The opening operator is the combination of erosion ⊖ and dilation ⊕: **B** = (**I** ⊖ **Φ**) 0 **Φ** (Van Den Boomgaard and Van Balen, 1992), where the morphological erosion returns minimum value (**I** ⊖ **Φ**)(*x, y*) = min_(*s,t*)∈**Φ**_{**I**(*x* + *s, y* + *t*)} and dilation returns maximum value (**I** ⊕ **Φ**)(*x, y*) = max_(*s,t*)∈**Φ**_{**I**(*x* + *s, y* + *t*)} within the same structuring window at each point. In practice, the choice of the structuring element should be comparable to the overall size of the neurons in the imaging field. We choose the structuring element to be of disk shape with a default size of 9 (pixels). The foreground is then computed as: **I^f^** = **I** – **B**. After the dynamic background removal, **I^f^** contains only neuronal information with minimal background noise corruption.

### MOVEMENT CORRECTION

The movement correction module performs a hierarchical registration framework over the frames of the imaging video. The neural enhanced video is first scored with a measuring metric on the relative displacement between two neighboring frames. The video is then segmented into stable or nonstable sections based on the score, followed by three levels of registration: the intra-stable-section, inter-stable-section and nonstable-section registration.

#### Scoring Metric

We track features containing corner information of every two neighboring frames using the KLT tracker, and then calculate the average displacement of these features. The KLT tracker first selects good features to track, where the feature points contains enough information on both directions within the neighborhood *p*. The problem can be formulated as ∇*p* = 0 over the neighborhood. A good feature point is selected if the smaller eigenvalue *σ_min_* ≥ *σ*_0_ and the *condition number* 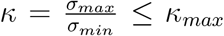. Then the algorithm tracks the feature points based on Newton-Raphson minimization on the normal equation:

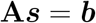

where

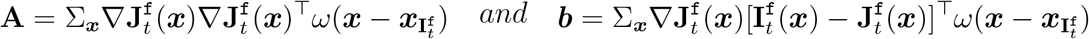

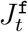 and 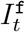 are the moving and fixed frame of the *t*th registration pair after neural enhancing respectively (in this case, 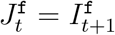), and *ω*(*x*) is the window of the neighborhood. For the scoring, we choose 31 pixels as the default window size of the neighborhood. To segment the frames into stable/nonstable sections, we compute the minimum value between the 3 times of median absolute deviation (MAD) above median and upper limit threshold (0.5 pixel as default) to be the threshold of segmentation. Any frame whose score is above the threshold is considered as one belonging to some stable section. The frames that are connected together are labeled as stable sections while the remaining connected components are nonstable ones.

#### Intra-stable-section Registration

Displacements with small magnitude or consistent directions can be approximated as translational displacement. Therefore, after the segmentation, movements within the stable sections can be considered translational. Within each stable section, an efficient displacement matching method with subpixel resolution is used to register the neighboring frames. Here we use a similar Lucas-Kanade tracker to track the center of the moving frames with a sufficiently large window (80% of the frame size) between two neighboring frames. However, when frame number within a section grows, the registration error may cumulate for later frames. Therefore, in practice we update the fixed frame as reference every second and all moving frames within that period are aligned to the fixed frame. This is acceptable due to the slow calcium dynamics so that the imaging field remains essentially similar within a second.

#### Inter-stable-section Registration

The imaging field is stable within each stable section after the intra-section registration. However, the imaging fields of different stable sections are not necessarily similar. Therefore, we need to align different stable sections without specific assumptions of the movement type. We first extract the overall information of each stable section by averaging the frames with significant neuronal activation, we then sort these sectional images 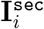 according to their similarity. To register the ith sectional image based on the sorted similarity, a reference image is generated by calculating the weights 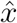 using the least square regression between the registered frames and the current sectional image:

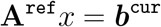

where

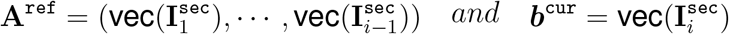

The reference image is then reshaped into two dimensional matrix 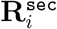 from 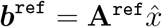. Once we have the moving and fixed sectional images, we apply rigid preregistration to the moving image using similar KLT tracker. To finally correct for the nonrigid deformation, we apply the diffeomorphic Log-Demons to the preregistered image 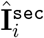 based on the reference 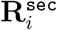. In general, the diffeomorphic Log-Demons is a non-parametric registration method, which aims at finding the displacement of all pixels by minimizing the global energy:

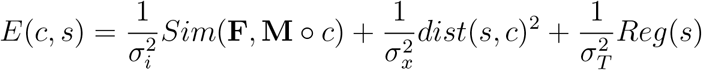

It consists of the similarity, correspondence and regularization items, where **F** and **M** are fixed and moving image respectively, and *σ_i_, σ_x_* and *σ_T_* account for similarity, spatial uncertainty on the correspondence and regularization level. In specific, Log-Demons represents everything in log domain. It spells out two regularizing terms, fluid and diffusion, and uses a Lie group structure to impose diffeomorphism. In our implementation, we choose *σ_i_* = *σ_x_* = 1, and *σ_fiuid_* = *σ_diffusion_* = 3. Because all the frames within the same stable section share the same imaging field, the deformation field of the current stable section is then applied to warp all the frames. After registering all the stable sections iteratively, all the stable frames are efficiently aligned.

#### Nonstable-section Registration

At the third level, only nonstable sections remain to be aligned. We use a similar approach as in the inter-section registration, where we first preregister the current frame to the reference image, and then apply Log-Demons to finely align the frame. Similarly, the reference image is computed based on some of the registered frames. For the ith nonstable section, this set of frames includes the last frame of the *i*th stable section, the first frame of the *i* + 1th stable section, and the frames that have already been registered in the *i*th nonstable section.

### SEEDS-CLEANSED SIGNAL EXTRACTION

We generate and cleanse the potential seeds of ROIs in this section, and create the initial spatial regions and the time series of ROIs for later refinement of the spatiotemporal signals.

#### Over-complete Seeds Initialization

To generate the initial set containing centers of all potential ROIs, we construct a *randomized max pooling* process to create an over-complete set of seeds. This process first randomly selects a portion of frames 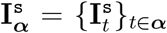, ***α*** ⊆ {1,…, *T*}, where ***α*** is a randomized subset of frame number. We then compute the max-projection map across selected frames 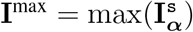, and further detect all the local max points on this map as 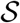. We repeat the above procedures multiple times, and collect the union of each map 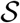 as the final over-complete set of neural seeds 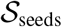. Our randomized max pooling can improve the true positives compared with max-pooling over the entire video, because a real seed can be buried in the uneven florescence of an ROI over a long period but can most likely be discovered in a small temporal vicinity. To exclude false positive seeds from the set, we propose the following two-stage algorithm for seeds refinement.

#### Seeds Refinement with GMM

The temporal properties of ROIs and non-ROIs can be very different. In one aspect, the ROIs often have prominent peak-valley difference *d*, while non-ROIs tend to be less spiky. We assume 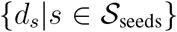 are generated from a mixture of two Gaussians 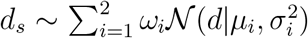, where *μ_i_*, 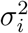, *ω_i_* indicate the mean, variance, mixture proportion of the *i*th Gaussian component. Therefore, we can cluster the seeds based on their probabilities of belonging to each component, and only consider those having higher probabilities to the Gaussian with a larger mean as positive seeds.

#### Seeds Refinement with LSTM

To select true ROI seeds from the over-complete set, we need to classify the seeds based on their patterns of neuronal calcium dynamics. We propose to employ Recurrent Neural Networks (RNNs) (LeCun et al., 2015) for calcium signal sequence classification. We offline trained the RNNs with a separate training dataset, composed of both positive and negative sequence chunks of length *T*_0_ = 100, obtained with 10Hz frame rate. The training data for the RNN module were selected from a separate real dataset. The labels were manually selected by experienced neuroscientists with a conservative standard: the positive trials were the ones with most obvious calcium dynamics that were aligned to the peak of the spike, whereas the negative trials were randomly selected from the rest of the data. The training dataset contained 1000 positive and 1000 negative labels, and both the validation and the test set contained 600 balanced trials. Specifically, consider training data 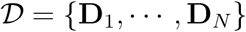, where 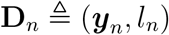, with input sequence ***y**_n_* and output label *l_n_* ∈ {0,1}. Our goal is to learn model parameters ***θ*** to best characterize the mapping from ***y**_n_* to *l_n_* with likelihood 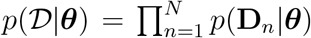. In our setting for sequence classification, the input is a sequence, ***y*** = {*y*_1_,…, *y*_*T*_0__}, where *y_t_* is the input data at time *t*. There is a corresponding hidden state vector 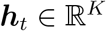 at each time *t*, obtained by recursively applying the *transition function **h**_t_* = *g*(***h***_*t*–1_, ***y**_t_*; **W**, **U**). **W** is *encoding weights*, and **U** is *recurrent weights*. The ouput *c* for our classification is defined as the corresponding *decoding function p*(*c*|***h***_*T*_0__; **V**) = *σ*(**V*h***_*T*_0__), where *σ*(·) denotes the *logistic sigmoid function*, and **V** is *decoding weights*.

The transition function *g*(·) can be implemented with a *gated* activation function, such as LSTM (Hochreiter and Schmidhuber, 1997) or a Gated Recurrent Unit (GRU) (Cho et al., 2014). Both LSTM and GRU have been proposed to address the issue of learning long-term sequential dependencies. Each LSTM unit has a cell containing a state c_*t*_ at time *t*. This cell can be viewed as a memory unit. Reading or writing the memory unit is controlled through sigmoid gates: input gate i_*t*_, forget gate f_*t*_, and output gate o_*t*_. The hidden units ***h**_t_* are updated as follows:

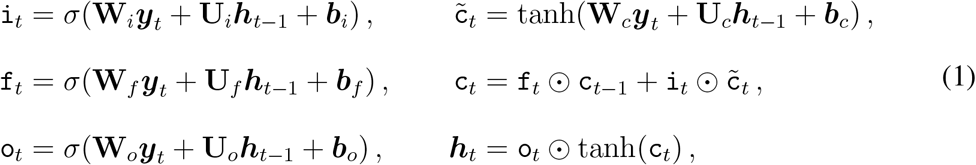

where ⊙ represents the element-wise matrix multiplication operator. Note that the training of RNNs is completed off-line, only the efficient testing stage is performed for seeds refinement. In the testing stage, given an input 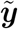 (with missing label 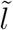), the estimate for the output is 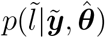, where 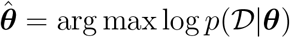. In our practical application, the testing sequence 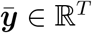 is often of length *T* > *T*_0_. We first convert it into abag of subsequences 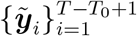, with a sliding window of width *T*_0_ and moving step size 1. The well trained LSTMs are then used to label the subsequences. We consider 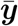 as positive if at least one subsequence in its bag is classified as “calcium spike” (*l* = 1), otherwise negative. Intuitively, this means that we only care about the patterns of neural spikes, regardless of their temporal positions.

#### Seeds Merging

Once the set of seeds is cleansed, there is still a low possibility of identifying multiple seeds within a single ROI. Therefore, we merge all these redundant seeds by computing the temporal similarity of seeds within their neighborhood, and preserving the one with maximum intensity. Specifically, we compute the similarity based on phase-locking information (Hahn et al., 2006). For a sequence, we extract the instantaneous phase dynamics using Hilbert transform, and only consider the subsequences containing prominent peaks, because all the pixels within the same ROI are highly correlated only during the calcium spiking periods, but not necessarily during baseline period. After seeds merging, we obtain *K* seeds, as the number of ROIs in our MIN1PIPE.

#### Spatial and Temporal Initialization

The time series of the *k*th seed is used as initial guess of temporal signal ***ŝ**_k_*. The spatial map of the corresponding ROI, 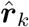, is estimated by pooling neighbor pixels with temporal similarity above a threshold. Because of our seeds generating and cleansing approach, the spatiotemporal initialization in our method can be readily parallelized. Then we fine-tune the corresponding spatial and temporal guess, by applying semi-NMF:

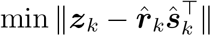

where ***z**_k_* ⊂ **I^s^** is a region containing the *k*th ROI. The spatial map and the temporal signal are updated respectively:

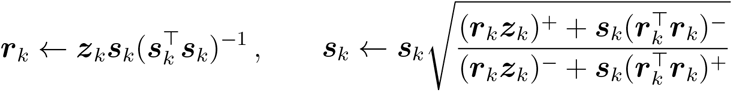

where 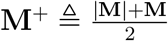 and 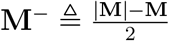 for any matrix **M** (Ding et al., 2010). In total, we now have *K* ROIs **S**^0^ = [***s***_1_, ···, ***s**_K_*], and temporal signals **R**^0^ = [***r***_1_, ···, ***r**_K_*]. Similarly, the background parameters ***b***^0^, ***f***^0^ are also estimated by the same semi-NMF procedure, using 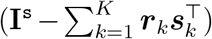. **S**^0^, **R**^0^, ***b***^0^, ***f***^0^ are then used as initializations of the spatiotemporal signal refinement.

#### Spatiotemporal Signal Refinement

We perform the iterative spatial and temporal optimizations to update the spatial footprints of individual ROIs, and the temporal traces with deconvolved spike trains, as proposed in CNMF. Specifically, for the neural enhanced video **X^f^**, the method decomposes **X^f^** into a spatial dictionary 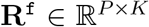 representing individual ROIs, and corresponding temporal dynamics matrix 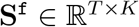, in addition to the background **B^f^** and the background dynamics **E**:

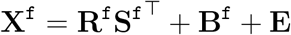

Similarly, with the rank-1 assumption on **B^f^**, CNMF decomposes the background **B^f^** = ***b*^f^f**^**f**^T^^, where 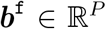 and 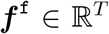. Meanwhile, **S^f^** is also correlated with underlying action potential events: **A^f^** = **S**^**f**^T^^**G^f^**, where 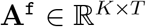, and 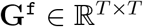 is the coefficient matrix of the low-order autoregression. The variables **R^f^**, **S^f^**, **b^f^**, ***f*^f^** are therefore estimated via iteratively alternating between the following two steps [26].

Estimating Spatial Variables Given the estimates of temporal variables **S**^**f**(*ℓ*–1)^ and ***f*^f^**^(*ℓ*–1)^ from the last iteration, the spatial parameters can be updated by solving the problem:

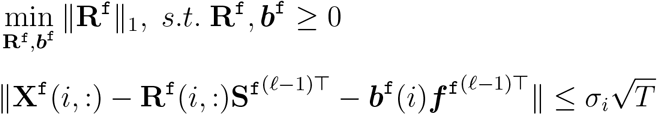

where **X^f^**(*i*,:) is the *i*th row of the matrix **X^f^, *b*^f^** (*i*,:) is the ith element of the vector. 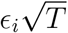 is the empirically selected noise residual constraint of the corresponding pixel. This is essentially a basis pursuit denoising problem, and it is solved by the least angular regression (Efron et al., 2004) in implementation.

#### Estimating Temporal Variables

Given the estimates of spatial variables **R^f^**, ***b*^f^** and temporal parameters **S**^**f**(*ℓ*–1)^, ***f***^**f**^(*ℓ*–1)^^ from the last iteration, the temporal parameters can be updated by solving the problem:

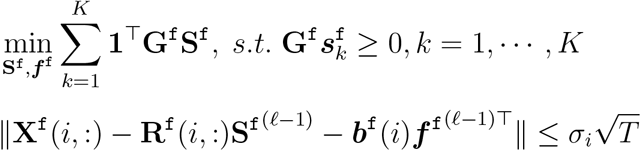

Unlike in CNMF, where the spatial footprints are serially updated and subtracted from the preceding residuals, we extract spatial footprints from the original data that does not depend on preceding iterations. Therefore, the information loss/duplication is reduced and the optimization procedures can be parallelized in our method.

#### Data availability

The data that support the findings of this study are available from the corresponding authors upon request.

#### Code availability

The codes will be freely available upon publication, and the beta version is available for reviewers upon request.

